# NanoPyx: super-fast bioimage analysis powered by adaptive machine learning

**DOI:** 10.1101/2023.08.13.553080

**Authors:** Bruno M. Saraiva, Inês M. Cunha, António D. Brito, Gautier Follain, Raquel Portela, Robert Haase, Pedro M. Pereira, Guillaume Jacquemet, Ricardo Henriques

**Author notes:** These authors contributed equally to this work (Bruno M. Saraiva, Inês M. Cunha, António D. Brito).

## Abstract

To overcome the challenges posed by large and complex microscopy datasets, we have developed NanoPyx, an adaptive bioimage analysis framework designed for high-speed processing. At the core of NanoPyx is the Liquid Engine, an agent-based machine-learning system that predicts acceleration strategies for image analysis tasks. Unlike traditional single-algorithm methods, the Liquid Engine generates multiple CPU and GPU code variations using a meta-programming system, creating a competitive environment where different algorithms are benchmarked against each other to achieve optimal performance under the user”s computational environment. In initial experiments focusing on super-resolution analysis methods, the Liquid Engine demonstrated an over 10-fold computational speed improvement by accurately predicting the ideal scenarios to switch between algorithmic implementations. NanoPyx is accessible to users through a Python library, code-free Jupyter notebooks, and a napari plugin, making it suitable for individuals regardless of their coding proficiency. Furthermore, the optimisation principles embodied by the Liquid Engine have broader implications, extending their applicability to various high-performance computing fields.

## Introduction

Super-resolution microscopy has revolutionised cell biology by enabling fluorescence imaging at an unprecedented resolution (4–7). However, the data collected from super-resolution experiments requires specific analytical procedures, such as drift correction, channel alignment, resolution enhancement, and quantifying data quality and resolution. Many of these procedures use open-source image analysis software, particularly ImageJ (8) or Fiji (9); and associated plugins such as ThunderSTORM (10), Picasso (11), FairSIM (12), Fourier Ring Correlations (FRC) (13), and Decorrelation Analysis (14). The computational performance of these methodologies bears significant implications for processing time and becomes especially salient given the increasing need for high-performance computing in bioimage analysis.

Computational performance has emerged as a significant bottleneck with the expanding adoption of super-resolution microscopy and the consequent upscaling of datasets (number, size, and complexity). This has highlighted the need for a shift towards a more performance-centric approach in managing increasingly extensive datasets and addressing the limitations currently experienced in (super-resolution) microscopy methodologies.

Here, we introduce NanoPyx, a high-performance and adaptive bioimage analysis framework. NanoPyx is not only a Python library but also provides code-free Jupyter notebooks (15) and a napari (16) plugin. At the core of NanoPyx is the Liquid Engine, an agent-based machine-learning system that predicts acceleration strategies for image analysis tasks. To enhance its image analysis capabilities, the Liquid Engine uses multiple variations (here referred to as implementations) of the same algorithm to perform a specific task. Although these implementations provide numerically identical output for the same input, their computational performance differs by exploiting different computational strategies. OpenMP is used for parallelising the code at the CPU level, while OpenCL (17) is used at the GPU level. The Liquid Engine then employs a specially created machine-learning system to predict the optimal combination of implementations based on the user”s specific computational environment. This creates a competitive setting wherein various algorithm implementations are benchmarked against each other, to achieve the highest performance.

One of the strengths of NanoPyx is its modular design, which enables users to easily include NanoPyx as part of new methods and algorithms. In its initial iteration, NanoPyx enhances and expands the super-resolution analysis methods previously included in the NanoJ library while introducing a new Python implementation of the Decorrelation Analysis method (14). NanoJ is an extensive suite of ImageJ plugins custom-built for the domain of super-resolution microscopy. Notable methods exploiting NanoJ (1) include NanoJ-SRRF (2) and NanoJ-eSRRF (18) which generate super-resolution reconstructions from diffraction-limited image sequences; and NanoJ-SQUIRREL (3) which provides image quality and resolution analysis. Bringing the adaptability of the Liquid Engine into these methods allows NanoPyx to overcome many limitations of NanoJ and other modern bioimage analysis packages. By providing a flexible framework we can assure accessibility of both new and old image analysis pipelines regardless of the user hardware, with further performance enhancement. Furthermore, we can leverage this flexibility and use it in conjunction with other Python libraries and tools. This is particularly valuable as many methods increasingly rely on Python-based deep learning techniques, making the use of current analysis frameworks (such as NanoJ (1)) prohibitive in specific scenarios.

As part of NanoPyx, users can access critical features, including drift correction (1), channel registration (1), SRRF (2) and eSRRF (18), error map calculation as per NanoJ-SQUIRREL (3), Fourier Ring Correlation (FRC) (13), and Image Decorrelation Analysis (14) (1). To make NanoPyx accessible to users with different levels of coding expertise, we offer it through three separate avenues -as a Python library for developers with the skills to create their workflow scripts, as Jupyter Notebooks (15) that can be executed either on a local machine or on a cloud-based service like Google Colaboratory, and as a plugin for napari (16), a Python based image viewer, for users without programming experience. By distributing NanoPyx in this manner, we can cater to the needs of a wider audience, ensuring users of varying coding expertise have easy access and can effectively utilise NanoPyx for their bioimage analysis needs.

## Results

### NanoPyx’s Liquid Engine

Pure Python code often runs on a single CPU core, impacting the performance and speed of Python frameworks. Alternative solutions, such as Cython (20), PyOpenCL (21) and Numba (22), permit parallelisation of the CPU and GPU, while enabling a considerable computational acceleration (Supplementary Note 1). However, identifying the swiftest implementation depends substantially on the input data and available hardware. For instance, Figure 2 presents a case where for smaller inputs, employing threaded CPU processing completes an interpolation task over 10x faster than a GPU. However, the situation reverses with increasing input size, where GPU-based processing reveals itself as the more efficient alternative. In Supplementary Figure S1, a comprehensive analysis is conducted to contrast the execution times of OpenCL with alternative implemented run time methodologies using diverse hardware configurations, including Google Colaboratory. The obtained results not only reaffirm the identified correlation between the faster implementation and input data size, but also elucidate the hardware-dependent breakpoints at which a given implementation surpasses the performance of its alternatives. Supplementary Figures S2, S3 and S4 further elucidate these observations by illustrating run times for various implementations across distinct input datasets and parameters on two contrasting hardware setups. The benchmark used features a 2D convolution with varying kernel sizes. While the professional workstation results align with expectations -OpenCL implementation was markedly faster as input image size and kernel size increased. However, this was not mirrored on a laptop device (Supplementary Figure S2). Laptop performance showed that while larger kernel sizes boosted OpenCL”s relative efficiency against CPU threading, expanding image size beyond a certain threshold made the parallelised CPU approach faster again. Notably, this outcome likely ties to the test laptop (MacBook Air M1) lacking a dedicated GPU, demonstrating how closely run times are tied to specific user hardware. This apparent disparity in results underlines how reliance on one implementation can prove restrictive; for instance, choosing OpenCL implementation for lower-sized images could escalate the run time by up to 300 times compared to CPU processing. Similarly, threaded CPU processing for larger-sized images performed up to 10x slower than GPU processing on professional workstations. Collectively these findings stress the importance of having an adaptable system that selects the optimum implementation based not just on data inputs but also considering unique user hardware configurations.

**Fig. 1.**
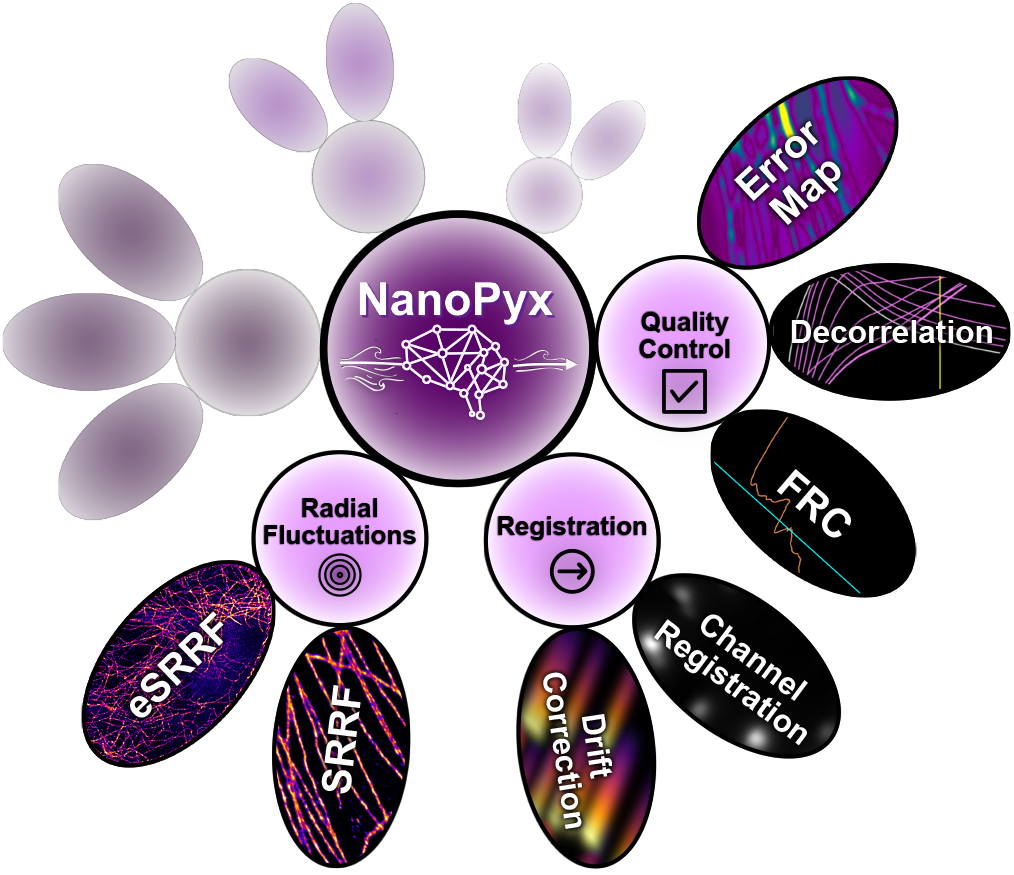
Schematic representation of the NanoPyx framework. NanoPyx is a Python framework for super-resolution microscopy images. It uses the Liquid Engine for self-tuning high performance. Currently, NanoPyx offers methods for Image Registration (1), Radial Fluctuations (2), and Quality Control (3) categories.

**Fig. 2.**
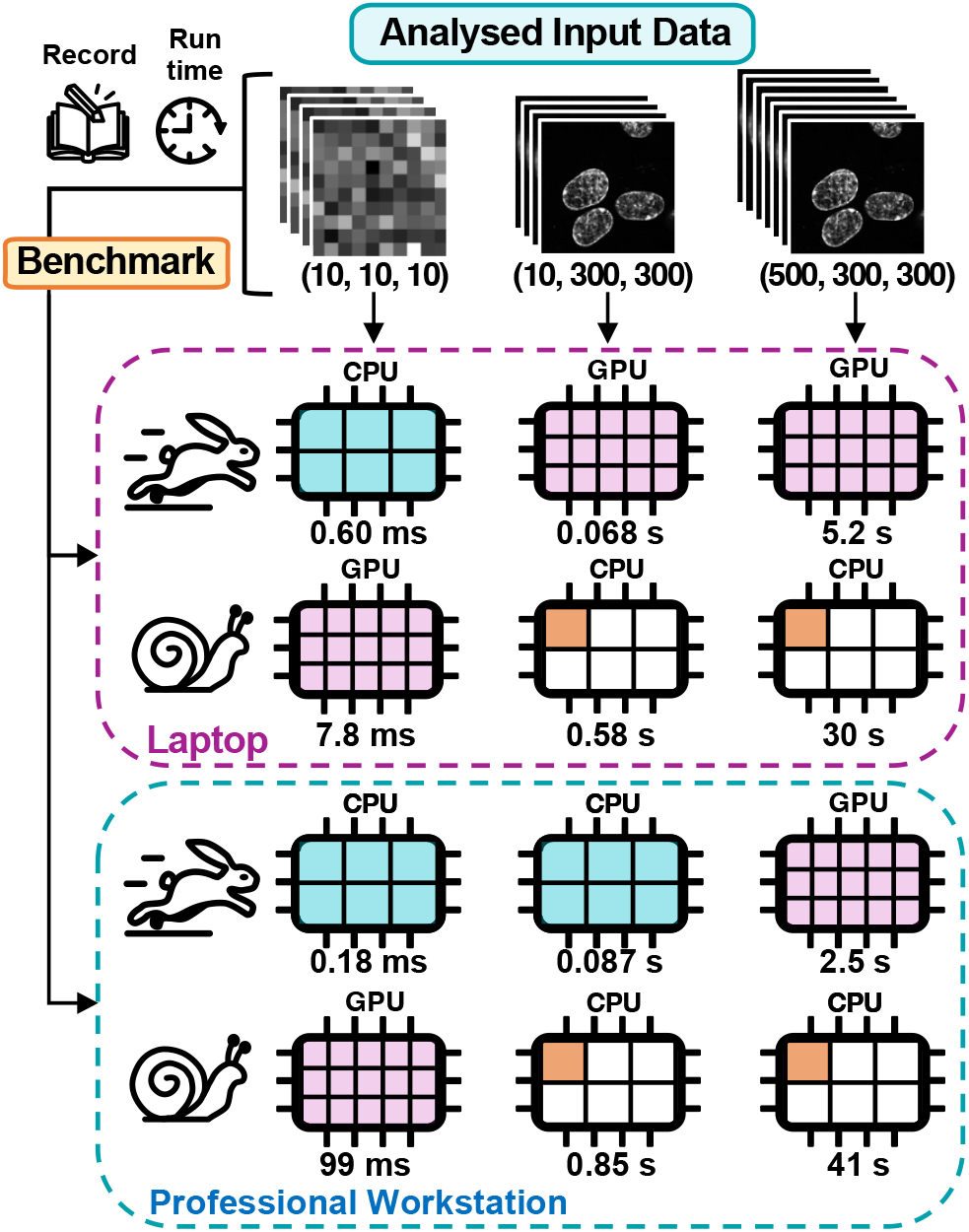
Comparative run times of multiple implementations of an algorithm, ran on either a consumer-grade laptop or a professional workstation. The fastest (rabbit) and slowest (snail) implementations depend on the shape of the input data and the user device. The underlying task carried out is a 5x frame-wise bicubic (19) upscaling. Different implementations are represented as processing chips with different colours. The implementation of both threaded (blue chip) and unthreaded (white chip with orange core) CPU uses optimised Cython code, while the implementation for GPUs (pink chip) is done through OpenCL.

To address this, we have developed the Liquid Engine. This machine learning-based system manages multiple tasks by exploiting various device components and selecting the most efficient implementation based on input data (Supplementary Note 2). The Liquid Engine can significantly enhance computational speed for tasks involving input data of varying sizes. It achieves this by predicting when to switch between algorithmic implementations, as depicted in Supplementary Figures S1 and S2, showing the capacity for a 10x acceleration. The black dotted line in Figure S2 indicates when the switch between implementations occurs. The Liquid Engine features three main components: meta-programming tools for multi-hardware implementation (called tag2tag and c2cl, see Supplementary Note 3); an automatic benchmarking system for different implementations; and a supervisor machine learning-based agent that determines the ideal combination of implementations to maximise performance (Figure 3).

**Fig. 3.**
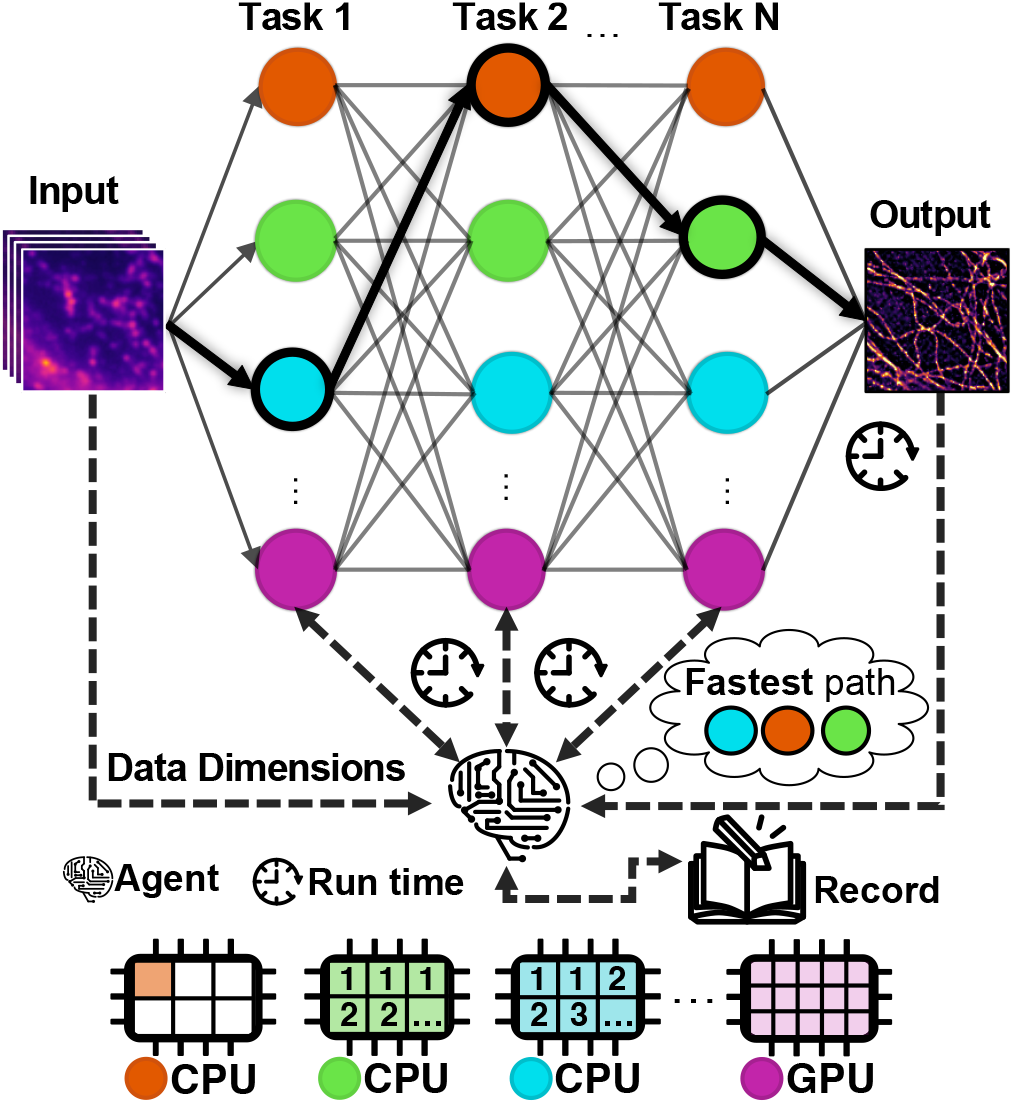
NanoPyx achieves optimal performance by exploiting the Liquid Engine self-optimisation capabilities. The image analysis workflows of NanoPyx are built on top of the Liquid Engine, which automatically benchmarks implementations of all tasks in the specific workflow. The Liquid Engine keeps a historical record of the run times of each task and the shape of the used input, allowing a machine-learning-based agent to select the fastest combination of implementations. In the case of an unexpected delay, the agent dynamically adjusts the preferred implementations to ensure optimal performance.

The tag2tag tool enables developers to generate multiple implementations of the same algorithm written as C or Python code snippets. Effectively, tag2tag transcribes these snippets into single-threaded and multi-threaded versions of the code, generally then called by Cython (20). In addition, the c2cl tool auto-generates GPU-based implementations based on C code snippets, using OpenCL (17), called via Py-OpenCL (21). The Liquid Engine also supports Numba (22) as an alternative performance-boosting option for Python code snippets. The implementation of the Liquid engine in NanoPyx adapts to the user’s hardware, selecting the fastest implementation available to each user and ensuring optimal computational speeds. In the cases where a user does not have access to one of the implementations, it will ignore that implementation and pick the fastest from the remaining ones, guaranteeing that users will always be able to process their images.

### Liquid Engine’s adaptive optimisation

NanoPyx’s Liquid Engine independently identifies ideal implementation combinations for specific workflows keeping in view device-dependent performance variations (Figure 2 and Supplementary Figure S1 and S2). Through automatic benchmarking of each implementation, the Liquid Engine keeps an historic record of runtimes for each implementation. Whenever a workflow is scheduled to be run, the supervisor agent is responsible to select the optimal implementation based on the previous recorded run times. The agent can adapt to unexpected delays in any implementation (Supplementary Figure S5). In case a severe delay is detected, reaching a level where it could potentially lead to a different implementation becoming faster, the agent predicts if the optimal implementation has changed. For that, the agent predicts the likelihood of the delay being repeated in the future and then assigns a probability for each implementation that depends on an estimation of the expected run time that each one might take. For instance, if the fastest implementation for a method uses OpenCL (17) and the GPU is under heavy load, resulting in an abnormally prolonged run time that is longer than the second fastest, the agent activates its delay management (Supplementary Figure S6). All available implementations are now assigned a probability that is a function of their expected run time, as given by the average values measured in the past. The expected run time for the delayed implementation is adjusted based upon the probability that the delay is maintained and the magnitude of the measured delay itself. Therefore, in this example, the probability the agent chooses to run using OpenCL is low, especially if the delay is continuously maintained. However, the delayed implementation should always have a bigger than zero probability to be chosen. Due to this probabilistic approach, the agent will still select and use the delayed implementation from time to time. This ensures that it can detect when the delay is over. Once it detects that the delay is over, the agent goes back to selecting implementations based only on the fastest average run time. In the example of Supplementary Figure S6, the Liquid Engine was able to detect an artificial delay that slowed down the OpenCL implementation. During the delay, the OpenCL implementation was used less times, but stochasticity allowed the Engine to detect the end of the delay. In this example, over the course of several sequential runs of the same method, we show that delay management improved the average run time by a factor of 1.8 for a 2D convolution and 1.5 for an eSRRF analysis (Supplementary Figure S6). Users can also manually initiate benchmarking, prompting the Liquid Engine to profile the execution of each implementation, using either multiple automatically generated data loads or using their own input, and identify the fastest one. This benchmarking is performed per task, allowing the Liquid Engine to adapt to the user’s hardware configuration and progressively optimise the chosen combination of implementations to reduce the total run time. The system analyses similar benchmarked examples from the user’s past data, using fuzzy logic (23) (see Supplementary Note 4) to identify the benchmarked example with the most similar input properties, utilising it as a baseline for the expected execution time. This system enables NanoPyx to immediately make adaptive decisions based on an initially limited set of benchmarked examples, progressively learning, and improving its performance as more data is processed.

### The NanoPyx Framework

NanoPyx is a comprehensive and extensible bioimage analysis framework providing wide-ranging methods which can cover an entire bioimage analysis microscopy workflows. In Figure 4, we showcase an example workflow where NanoPyx performs channel registration followed by drift correction to correct any chromatic aberration and drift that might occur during image acquisition. Once drift correction is completed, NanoPyx enables the generation of super-resolved reconstructions using the well-established Super-Resolution Radial Fluctuations (SRRF) (2) algorithm or its improved version eSRRF (18). Due to its focus on performance, NanoPyx can perform 2.5 times faster the same eSRRF (18) processing as NanoJ, with for the same input parameters. To ensure the fidelity of the reconstructions, NanoPyx incorporates rigorous quality control tools. The Error Map feature of NanoJ-SQUIRREL (3), implemented within NanoPyx, quantitatively assesses errors introduced by the reconstruction algorithm. The resolution-scaled error (RSE) (3) and resolution-scaled Pearson’s correlation coefficient (RSP) (3) are calculated by comparing the diffraction-limited image stack with the diffraction-limited equivalent of the reconstruction. Furthermore, NanoPyx incorporates Fourier Ring Correlation (FRC) (13) and Image Decorrelation Analysis (14) to determine image resolution. These various assessment methods enable a comprehensive quantitative evaluation of the resolution improvements achieved by the super-resolved reconstruction. NanoPyx also allows users to perform channel registration on the acquired or super-resolved images.

**Fig. 4.**
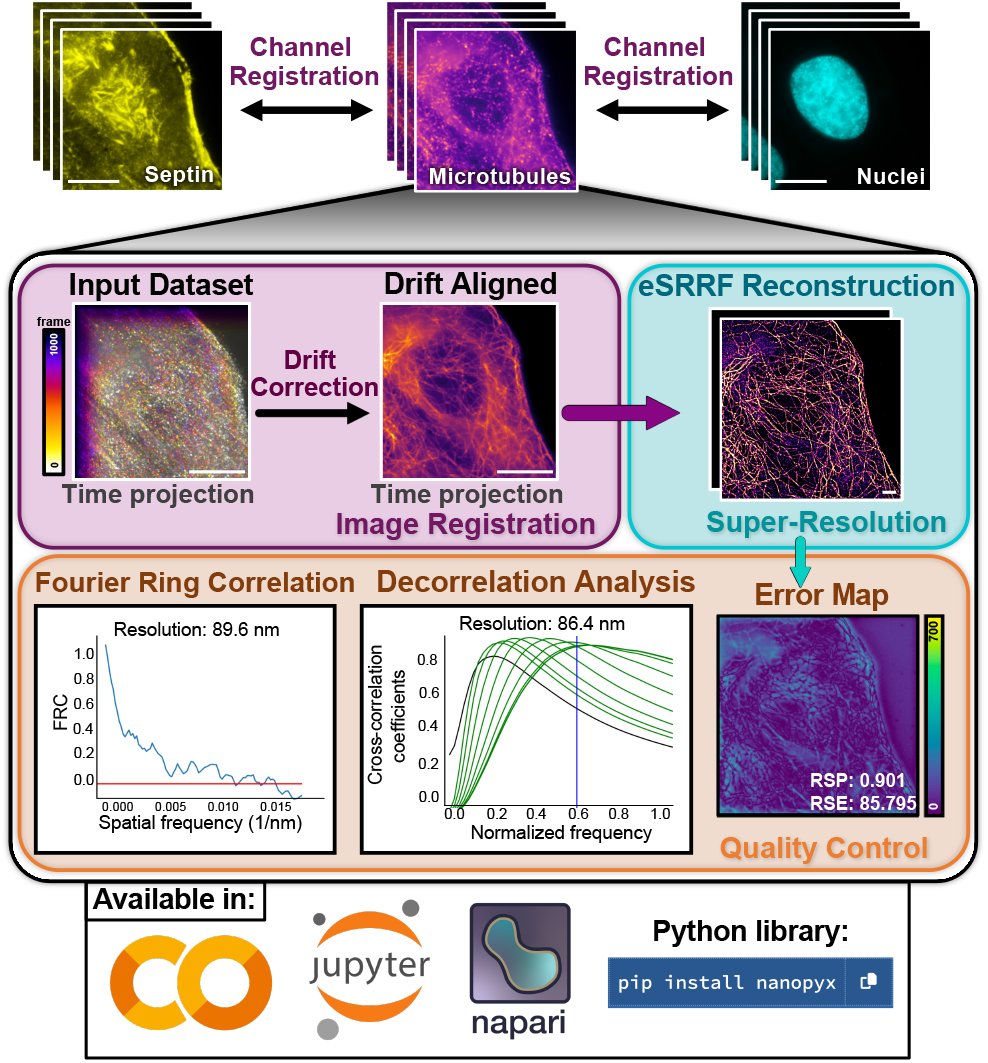
Microscopy image processing workflow using NanoPyx methods. NanoPyx implements several methods of super-resolution image generation and processing. Through NanoPyx, users can correct drift that occurred during image acquisition, generate a super-resolved image using enhanced radiality fluctuations (eSRRF)(2), assess the quality of the generated image using Fourier Ring Correlation (FRC)(13) or Image Decorrelation Analysis (14), perform artifact detection using the error map and then perform channel registration in multi-channel images. NanoPyx methods are made available as a Python library, a napari plugin, and Jupyter Notebooks that can be ran locally or through Google Colaboratory. Scale bars: 10 *µ*m.

### Distribution to end users

NanoPyx was developed with the primary objective of ensuring accessibility and ease of use for end users. To achieve this goal, we have made available three distinct interfaces through which users can interact with and utilise NanoPyx. Firstly, NanoPyx is accessible as a Python library (Figure 4), which can be conveniently accessed and installed via PyPI (Python Package Index) for stable releases or through our GitHub repository for the latest development versions. The Python library primarily targets developers seeking to incorporate NanoPyx’s methodologies into their workflows. Alongside the Python library we provide template files to help developers implement their own methods using the Liquid Engine. Secondly, we have provided Jupyter notebooks (15) through our GitHub repository (Figure 4 and Supplementary Figure S7). Each notebook offers separate implementations of each individual method. Users of these notebooks are not required to interact with any code directly: by sequentially executing cells, a graphical user interface (GUI) is generated (24, 25), enabling users to fine-tune the parameters for each step easily.

Consequently, these notebooks are specifically designed for users with limited coding expertise. Lastly, for users desiring a more interactive approach, we are concurrently developing a plugin for napari (16), a Python image viewer, granting access to all currently implemented NanoPyx methods (Figure 4 and Supplementary Figure S7). By offering these three diverse user interfaces, we ensure that NanoPyx can be readily utilised by users irrespective of their coding proficiency level. NanoPyx also offers a wide range of example datasets for the users to test and explore its capabilities (see Supplementary Note 5).

## Discussion and Future Perspectives

NanoPyx introduces a novel approach to optimise performance for bioimage analysis through its machine learning-powered Liquid Engine. This enables dynamic switching between implementations to maximise speed based on data and hardware. In initial experiments, NanoPyx achieves over 10x faster processing by selecting the optimal implementation. This has significant implications given the rapidly expanding scale of microscopy image datasets.

The Liquid Engine’s optimisation strategy diverges from traditional approaches of relying on single algorithms or implementations. Alternative Python tools like Transonic (26) and Dask (27) that accelerate workflows through just-in-time compilation or parallelism do not adapt implementations based on context. In contrast, the Liquid Engine continually benchmarks and collects runtime metrics to train its decision-making model. It is this tight feedback loop that empowers dynamic optimisation in NanoPyx. Furthermore, alternative libraries such as Transonic may, in the future, aid in generating further implementations for the Liquid Engine to optimise and explore, therefore cross-pollinating and benefiting both projects.

Beyond raw performance, NanoPyx also provides an accessible yet extensible framework covering key analysis steps for super-resolution data. Workflows integrate essential functions like drift correction, reconstruction, and resolution assessment. NanoPyx builds upon proven ImageJ plugins while migrating implementations to Python. The modular architecture simplifies integrating components into new or existing pipelines.

Future NanoPyx development will focus on several objectives. Incorporating advanced simulation tools will generate synthetic benchmarking data to further optimise and validate algorithms. Supporting emerging techniques like AI-assisted imaging and smart microscopes is a priority, as NanoPyx’s speed is critical for real-time processing during acquisition. Expanding the algorithm library will provide users with a more comprehensive toolkit. Careful benchmarking across diverse hardware will maximize performance from laptops to cloud platforms. Enhancing usability through graphical interfaces will improve accessibility. Fostering an open-source community will help drive continual innovation.

Looking ahead, a priority for NanoPyx is expanding support for emerging techniques like AI-assisted imaging and smart microscopes. As these methods involve processing data in real-time during acquisition, NanoPyx’s accelerated performance becomes critical. We plan to leverage the Liquid Engine’s auto-tuning capabilities to optimise pipelines on heterogeneous hardware. This could enable real-time AI to guide data collection, processing, and analytics. Additionally, we aim to incorporate more diverse reconstruction approaches beyond current methods like SRRF and eSRRF. We are actively testing prototypes to integrate deep learning super-resolution into NanoPyx in a modular way, allowing users a choice of classic or AI-based methods. The Liquid Engine’s ability to dynamically select optimised implementations of these complex neural networks will be crucial for acceptable run times. We are also pursuing integrations with other emerging reconstruction techniques from the imaging literature.

In summary, NanoPyx delivers adaptive performance optimisation to accelerate bioimage analysis while retaining modular design and easy adoption. The optimisation principles embodied in its Liquid Engine could extend to other scientific workloads requiring high computational efficiency. As data scales expand, NanoPyx offers researchers an actively improving platform to execute demanding microscopy workflows.

## Methods

### Mammalian cell culture

A549 cells (The European Collection of Authenticated Cell Cultures (ECACC)) were cultured in phenol red-free high-glucose, L-Glutamine containing Dulbecco’s modified Eagle’s medium (DMEM; Thermo Fisher Scientific) supplemented with 10% (v/v) fetal bovine serum (FBS; Sigma), 1% (v/v) penicillin/streptomycin (Thermo Fisher Scientific) at 37 °C in a 5% CO2 incubator.

### Sample preparation for microscopy

A549 cells were seeded on a glass bottom *µ*-slide 8 well (ibidi) at a 0.05 – 0.1 x 106 cells/cm2 density. After 24 h incubation at 37 °C in a 5% CO2 incubator, cells were washed once using phosphate-buffer saline (PBS) and fixed for 20 min at 23 °C using 4 % paraformaldehyde (PFA, in PBS). After fixation, cells were washed three times using PBS (5 min each time), quenched for 10 min using a solution of 300 mM Glycine (in PBS), and permeabilised using a solution of 0.2% Triton-X (in PBS) for 20 min at 23 °C. After three washes (5 min each) in washing buffer (0.05% Tween 20 in PBS), cells were blocked for 30 min in blocking buffer (5% BSA, 0.05% Tween-20 in PBS). Samples were then incubated with a mix of anti-α-tubulin (1 *µ*g/mL of clone DM1A, Sigma; 2 *µ*g/mL of clone 10D8, Biolegend; 2 *µ*g/mL of clone AA10, Biolegend) and anti-septin 7 (1 *µ*g/mL of #18991, IBL) antibodies for 16 h at 4 °C in blocking buffer. After three washes (5 min each) using the washing buffer, cells were incubated with an Alexa Fluor™ 647 conjugated goat anti-mouse IgG and an Alexa Fluor™ 555 conjugated goat anti-rabbit IgG (6 *µ*g/mL in blocking buffer) for 1 h at 23 °C. Cells were then washed thrice (5 min each) in washing buffer and once in 1X PBS for 10 min. Finally, cells were mounted with a GLOX-MEA buffer (50 mM Tris, 10 mM NaCl, pH 8.0, supplemented with 50 mM MEA, 10% [w/v] glucose, 0.5 mg/ml glucose oxidase, and 40 *µ*g/ml catalase).

### Image acquisition

Data acquisition was performed with the Nanoimager microscope (Oxford Nanoimaging; ONI) equipped with a 100 x oil-immersion objective (Olympus 100x NA 1.45) Imaging was performed using 405-nm, 488-nm, and 640-nm lasers for Hoechst-33342, AlexaFluor555 and AlexaFluor647 excitation, respectively. Fluorescence was detected using a sCMOS camera (ORCA Flash, 16 bit). For channel 0, a dichroic filter with the bands of 498-551 nm and 576-620 nm was used; for channel 1, a 665-705 nm dichroic filter was used. The sequential multicolor acquisition was performed for AlexaFluor647, AlexaFluor555 and Hoechst-33342. Using an EPI-fluorescence illumination, a pulse of high laser power (90%) of the 640-nm laser was used, and 10 000 frames were immediately acquired. Then, the sample was excited with the 488-nm laser (13.7% laser power), whereas 500 frames were acquired, followed by the 405-nm laser excitation (40% laser power), with an acquisition of another 500 frames. For all acquisitions, an exposure time of 10 ms was used.

### Liquid Engine’s agent

Run times of methods implemented in NanoPyx through the Liquid Engine are locally stored on users” computers and are associated with the used hardware. For OpenCL implementations, the agent also stores an identification of the device and is capable of detecting hardware changes. Whenever a method is run through the Liquid Engine, the overseeing agent splits the 50 most recent recorded runtimes into 2 halves: one with the 25 fastest run times (fast average) and one with the 25 lowest (slow average). Then it calculates the average of the 25 fastest run times for each implementation and selects the implementation with the lowest average runtime. Once the method finishes running, the agent checks whether there was a delay, which is defined by the last runtime being higher than the previously recorded average runtime of the fastest runs plus four times the standard deviation of the fastest runs (Equation 1). If a delay is detected (Supplementary Figure S5), the agent will also calculate the delay factor (DF, Equation 2) and will activate a probabilistic approach that stochastically selects which method to run. This is performed by using a Logistic Regression model to calculate the probability of the delay being present on the next run and adjusting the expected runtime of the delayed implementation according to Equation 3, while still using the fast average for all non-delayed implementations. Then, the agent picks which implementation to use based on probabilities assigned to each implementation using 1 over the squared normalized expected runtime (Equation 4). This stochastic approach ensures that the agent will still run the delayed implementation from time to time to check whether that delay is still present. The agent decides that the delay is over once the last runtime becomes smaller than the slow average minus the standard deviation of the slowest runs or higher than the fast average plus the standard deviation of the fastest runs (Equation 5). Once the delay is over, the agent will go back to selecting which implementation to use based only on the fast average of each implementation (Supplementary Figure S6).

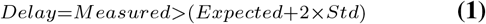

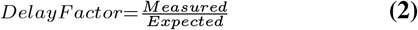

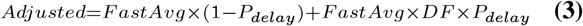

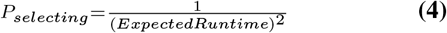

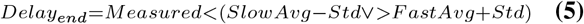

### Run times Benchmark

For the laptop benchmarks a MacBook Air M1 Pro with 16Gb of RAM and a 512Gb SSD was used. For the professional workstation, a custom-made desktop computer was used containing an Intel i9-13900K, a NVIDIA RTX 4090 with 24Gb of dedicated video memory, a 1TB SSD and 128Gb of DDR5 RAM. The first benchmark performed (Figure 2 and Supplementary Figure S1) was a 5 times up sampling of the input data, using a catmull-rom (19) interpolator. Benchmarks were performed on 3 different input images with shapes 10×10×10, 10, 10×300×300 and 500×300×300 (time-points, height, width). The second benchmarks (Supplementary Figure 2-4) were 2D convolutions using a kernel filled with 1s with varying sizes (1, 5, 9, 13, 17, 21) on images with varying size (100, 500, 1000, 2500, 5000, 7500, 10000, 15000 or 20000 pixels for both dimensions).

### NanoPyx comparison with NanoJ

Run times of eSRRF image processing were measured using a MacBook Air M1 with 16Gb of RAM and a 512Gb SSD. The parameters used for the analysis where the same for both NanoPyx and NanoJ: magnification – 5; radius – 1.5; sensitivity – 2; number of frames for SRRF – 1. The input image was a stack with 283 by 283 pixels and 10 000 frames. For the final image output an average reconstruction was performed.

### Availability

The NanoPyx Python library and the Jupyter Notebooks can be found in our Github repository: https://github.com/HenriquesLab/NanoPyx. The napari plugin implementing all NanoPyx methods can be found in a separate Github repository: https://github.com/HenriquesLab/napari-NanoPyx.

## ACKNOWLEDGEMENTS

We express our gratitude to the previous developers of the NanoJ framework, whose work inspired this study. Additionally, we extend thanks to Loic Royer and Juan Nunez-Iglesias for their invaluable feedback and guidance in preparing our work. R.H., P.M.P and R.P. acknowledge support from LS4FUTURE Associated Laboratory (LA/P/0087/2020). R.H., B.M.S. and I.M.C. acknowledge the support of the Gulbenkian Foundation (Fundação Calouste Gulbenkian), the European Research Council (ERC) under the European Union’s Horizon 2020 research and innovation programme (grant agreement No. 101001332), the European Union through the Horizon Europe program (AI4LIFE project with grant agreement 101057970-AI4LIFE, and RT-SuperES project with grant agreement 101099654-RT-SuperES), the European Molecular Biology Organization (EMBO) Installation Grant (EMBO-2020-IG-4734) and the Chan Zuckerberg Initiative Visual Proteomics Grant (vpi-0000000044 with DOI:10.37921/743590vtudfp). In addition, A.D.B acknowledges the FCT 2021.06849.BD fellowship. R.H. and B.M.S. also acknowledge that this project has been made possible in part by a grant from the Chan Zuckerberg Initiative DAF, an advised fund of Silicon Valley Community Foundations (Chan Zuckerberg Initiative Napari Plugin Foundations Grants Cycle 2, NP2-0000000085). P.M.P and R.P. acknowledge support from Fundação para a Ciência e Tecnologia (Portugal) project grant (PTDC/BIA-MIC/2422/2020) and the MOSTMICRO-ITQB RD Unit (UIDB/04612/2020, UIDP/04612/2020), P.M.P acknowledges support from La Caixa Junior Leader Fellowship (LCF/BQ/PI20/11760012) financed by “la Caixa” Foundation (ID 100010434) and by European Union’s Horizon 2020 research and innovation programme under the Marie Skłodowska-Curie grant agreement No 847648, and a from a Maratona da Saúde award. This study was supported by the Academy of Finland (338537 to G.J.), the Sigrid Juselius Foundation (to G.J.), the Cancer Society of Finland (Syöpäjärjestöt; to G.J.), and the Solutions for Health strategic funding to Åbo Akademi University (to G.J.). This research was supported by InFLAMES Flagship Programme of the Academy of Finland (decision number: 337531).

## AUTHOR CONTRIBUTIONS

B.M.S, P.M.P, G.J., R.He. conceived the study in its initial form; B.M.S., I.M.C., A.D.B., R.He. developed the NanoPyx framework with code contributions from R.Ha., G.J; B.M.S., I.M.C., A.D.B., R.He. designed the Liquid Engine optimization approach; B.M.S., I.M.C., A.D.B. implemented the Liquid Engine tools; G.F., R.P., P.M.P, G.J. provided samples, data, critical feedback, testing and guidance; B.M.S., I.M.C., A.D.B., G.F., G.J. performed experiments and analysis; B.M.S, P.M.P, G.J., R.He. acquired funding; B.M.S., P.M.P., R.Ha., G.J., R.He. supervised the work; B.M.S., I.M.C., A.D.B., G.J., R.He. wrote the manuscript with input from all authors.

## COMPETING FINANCIAL INTERESTS

The authors declare no conflict of interests.

## Supplementary Note 1: How computational acceleration for an algorithm implementation written with Cython, PyOpenCL and Numba is achieved

Python is an interpreted, high-level programming language that allows rapid development and easy code readability. However, Python’s flexibility and dynamic nature come at the cost of performance speed. Operations in pure Python are generally slower compared to compiled languages like C/C++. There are several methods to accelerate Python code by bypassing the Global Interpreter Lock (GIL) and compilation to machine code. Three popular approaches are Cython, PyOpenCL and Numba:

- Cython (1) is a static compiler that converts Python code into optimised C/C++ code that can be compiled into a Python extension module. It provides Python-like syntax while supporting calling C functions and declaring C types. Cython code runs significantly faster than Python because it bypasses the GIL to allow multi-threading and performs low-level optimisations like loop unrolling. One limitation is that Cython requires explicit type declarations, which removes some of Python’s dynamism. Overall, Cython can accelerate Python code 5-1400x faster (Supplementary Figure S1).
- PyOpenCL (2) allows Python code to execute parallel computations on Graphical Processing Units (GPUs) through the OpenCL framework. Computational tasks are offloaded to the GPU, which has thousands of tiny processing cores suited for data-parallel operations. PyOpenCL translates Python functions into OpenCL kernels that run efficiently on GPUs. This offers massive parallelism and speedup compared to Python limited by single-CPU execution. Despite PyOpenCL not requiring a physical GPU to run, the best and easiest performance improvement requires one, which can be a limiting factor for some users. Overall, PyOpenCL can accelerate some workloads up to 150x by harnessing GPU parallelism28 (Supplementary Figure S1).
- Numba (3) is a just-in-time (JIT) compiler that converts Python functions into optimised machine code using the LLVM compiler framework. It is designed to accelerate numerical and scientific workloads using NumPy arrays and math operations. Numba-compiled code avoids interpreter overhead and leverages vectorisation, loop-unrolling and parallel execution on multicore CPUs. But Numba has compilation overhead on first run. Overall, Numba can speed up math-heavy Python code by 100-400x by generating optimised machine code specialised for CPUs (Supplementary Figure S1).

In summary, all these three tools can significantly accelerate Python code by bypassing interpreter overhead and using efficient compilation, parallelism, and hardware optimisation. Cython translates Python to C/C++ code that can multi-thread and leverage CPU efficiency. PyOpenCL taps into massively parallel GPU hardware. Numba optimises machine code for numerical workloads on CPUs. Typical speedups depend on the methods used, nature of the code, size of the input data and hardware used.

## Supplementary Note 2: The machine-learning basis of the Liquid Engine

The NanoPyx Liquid Engine presents a straightforward machine-learning technique for performance self-tuning. To do so, it logs execution times of operations across multiple implementations like Python, Numba, Cython and OpenCL. These bench-marking times are used to train basic regression models to predict the occurrence of delays. If delays are detected the trained models are used at run time to estimate the magnitude and occurrence of a delay in a specific implementation. Over time, the benchmarking data is aggregated to refine the models continuously. This allows the Liquid Engine to “learn” the optimal implementations for a given platform, device, and data shape to maximise performance. It’s important to note that these principles do not use neural networks, rather focusing on looped data-driven optimisations. In summary, fundamental machine learning principles are applied in the engine itself for auto-tuning -the methods exposed to users are traditional image processing functions using optimised implementations under the hood.

## Supplementary Note 3: Meta-programming in the Liquid Engine

Meta-programming (4) is a programming technique where a program can manipulate or generate code during run time. In NanoPyx, meta-programming is used to generate optimised implementations of the same task automatically. The Liquid Engine uses two meta-programming tools: tag2tag and c2cl.

The tag2tag tool enables developers to generate multiple implementations of the same algorithm written as C or Python code snippets. Effectively, tag2tag transcribes these snippets into single-threaded and multi-threaded versions of the code, generally then called by Cython. Specifically, developers can write a single version of the code and delimit the “tag” to be propagated to the other implementations (e.g., a function). The tag2tag tool is able to read the content of the file, identify the tags, and store them in a dictionary, where the tag name is the key, and the associated code snippet is the value. After choosing the “tag”, in the same script, the developer can create a tag-copy, where they specify what part of the original tag should be replaced, and what to replace it for. tag2tag will identify the tag placeholders and apply the specified replace commands to the associated tag code. It then replaces the tag placeholder in the file with the modified tag code. This tool is intended for use with Python (.py), Cython (.pyx), and OpenCL (.cl) files. In practice, for most tasks implemented in the Liquid Engine, we used a single-threaded version of the code as the original tag, and created multi-threaded versions of the code by replacing the range in the for loops with prange. Additionally, we added implementations with different schedulers for the parallelisation. This allows the developer to easily create as many code variations as they find necessary. This approach streamlines the process of maintaining consistency across various code implementations, as altering the code in one version ensures that the modification is seamlessly and consistently applied to all other relevant versions. As a result, developers can effectively manage code updates and improvements, as these are propagated into the other implementations effortlessly, reducing redundancy and enhancing code maintainability.

Another meta-programming tool used in the Liquid Engine is the c2cl tool. c2cl is analogous to the tag2tag tool, but specifically designed to extract C functions and propagate them into .cl files, so the C functions can be used in OpenCL kernels. Manual conversion of these adaptations would be time-consuming and prone to errors, which is where c2cl comes in. The tool automates the process of porting C code to OpenCL by extracting reusable code blocks from the C functions, converting the C code into valid OpenCL kernels, and inserting the modified kernels back into the OpenCL file.

The Liquid Engine also supports Numba as an alternative performance-boosting option for Python code snippets. With all these implementations, NanoPyx can be run and used by users with diverse hardware configurations. Overall, the Liquid Engine extensively uses meta-programming techniques to avoid manual coding. Code generation and transformations are used to automatically create specialised implementations, resulting in a simple and flexible architecture.

## Supplementary Note 4: Fuzzy logic in the Liquid Engine

The Liquid Engine employs fuzzy logic (5) to match a specific function call to its most similar past benchmark.

The Liquid Engine adaptive nature is highly dependent on the existence of appropriate benchmarks for each implementation. As previously demonstrated, the time it takes for a particular image analysis task to execute is greatly influenced by the size of the input image and by the parameters chosen to perform the given task. To address this variability, benchmarks are stored separately for each unique set of parameters and data size. This approach allows the Liquid Engine to dynamically adjust its performance based on the specific conditions of each task, resulting in more accurate and optimised outcomes.

When the agent receives a request to execute a method using the Liquid Engine, it checks its records for historical run time data. However, when dealing with a new method executed with a unique combination of parameters, no existing benchmarks are available. To address this, the agent employs a strategy: it searches for the most similar set of parameters that has been previously used. This is because each set of parameters is associated with a run time score, which was saved along with the historical run time data in benchmark files. This score is determined by considering various factors, such as the dimensions of the input image and other relevant parameters. By finding the most similar parameters with known benchmark data, the Agent leverages this score to estimate and adapt the expected run time performance for the new method, even in the absence of specific benchmarks. This adaptive approach allows the Liquid Engine to intelligently adjust its behaviour and make informed decisions when executing methods with varying input conditions.

Finally, if no appropriate benchmarks exist for a specific run type, the score of the current parameters is compared to the score of all other benchmarks, and the benchmarks with the closest score are used.

## Supplementary Note 5: Example datasets in NanoPyx

NanoPyx provides users a wide range of example datasets. These datasets not only draw from our previous publications but also tap into publicly available datasets (6). These were integrated into the NanoPyx framework to facilitate testing and development, offering users an opportunity to gain hands-on experience and explore the capabilities of the library.

These include single-molecule localization microscopy data of Cos7 cells expressing Utrophin-GFP (from (7)); U2OS with microtubules labelled with AF647; Jurkat T cells expressing LifeAct-GFP (from (8)); Strucured Illumination Microscopy data of VACV A4 virions (from (9)); among others.

The management and loading of datasets within NanoPyx are orchestrated through a class specifically designed for data management, facilitating efficient access and use. The class provides functions to list datasets and retrieve their information, enabling users to effortlessly identify and choose relevant datasets. Most of the datasets are stored and accessed via Google Drive, and can be automatically downloaded as zip files. Once downloaded, these zip files can be effortlessly converted into numpy arrays, a process that seamlessly manages the complexities of image retrieval and manipulation.

In Jupyter or Colab notebooks, users can easily access the datasets through a user-friendly graphical interface (GUI) that has been developed to streamline the process. This interface empowers users to select datasets from a diverse array of options, all of which are thoughtfully named to provide clear context. This intuitive approach ensures efficient dataset selection and integration into analysis workflows.

## Supplementary figures

**Fig. S1.**
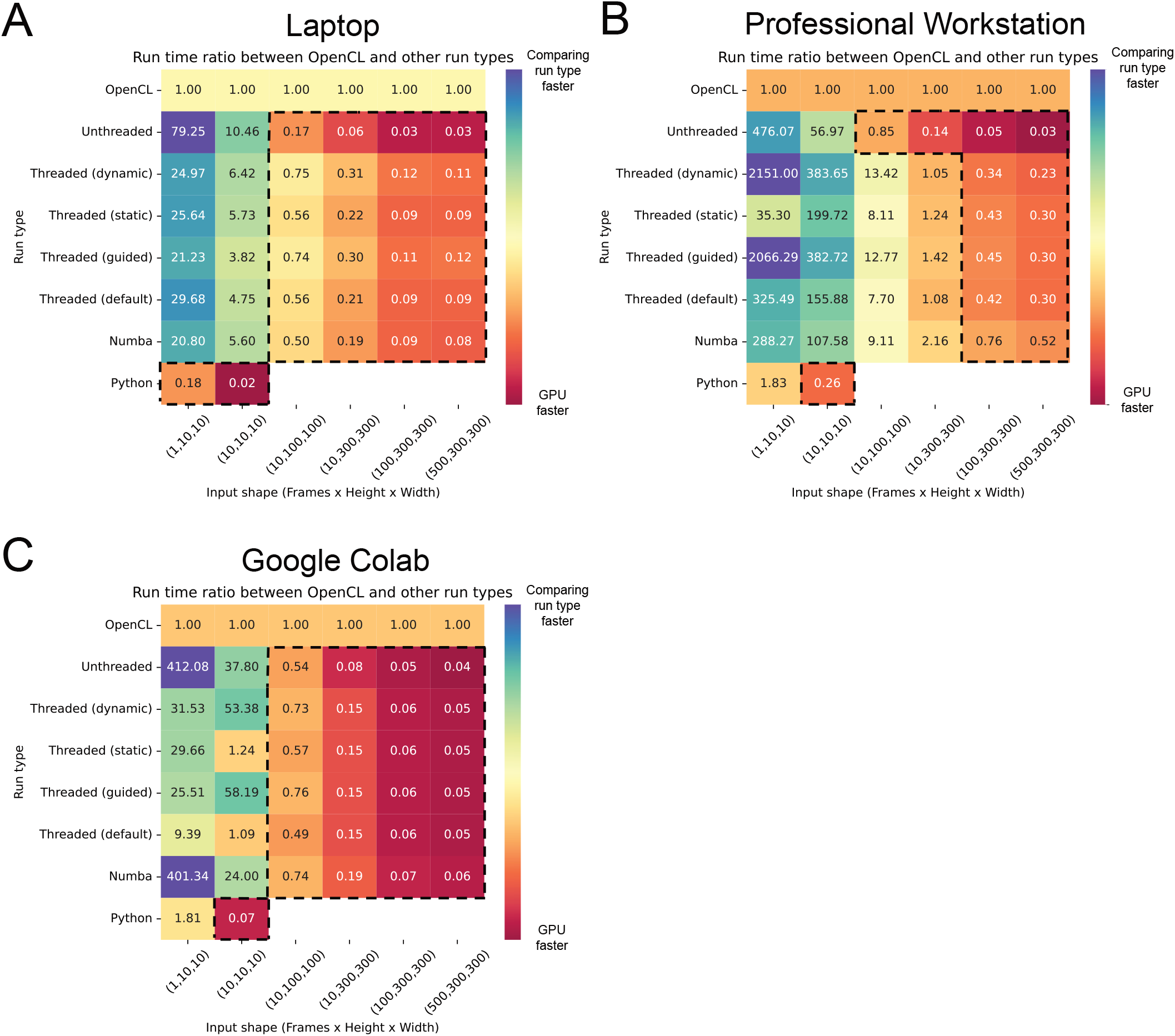
Ratio between the run times of OpenCL and other implemented run types. Run times of a 5x Catmull-rom22 interpolation were measured across multiple input data sizes using either a MacBook Air M1 (A), a Professional Workstation (B) or Google Collaboratory (C). Area within dashed lines correspond to kernel and image sizes where OpenCL is faster than other implementations.

**Fig. S2.**
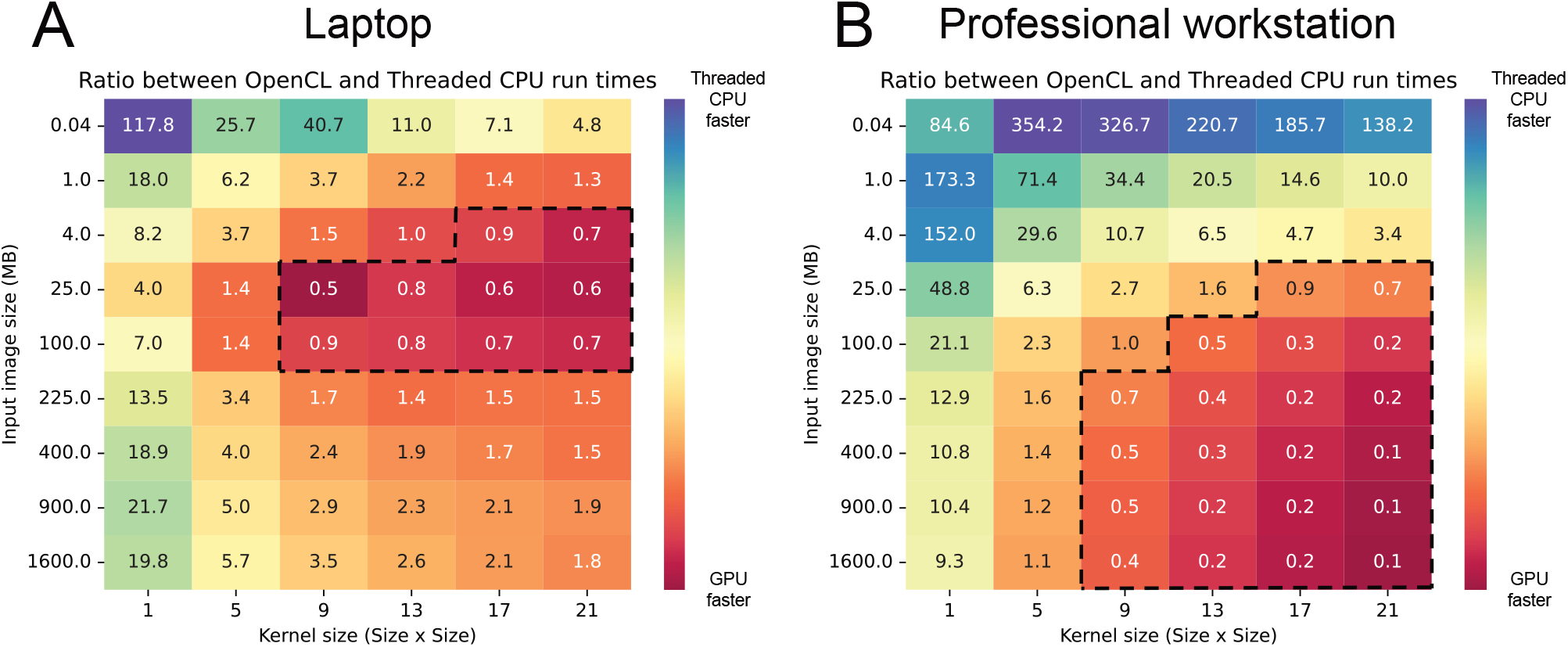
Ratio between the run times of a 2D convolution. Run times were measured across multiple input data sizes and kernel sizes using either a MacBook Air M1 (A) or a Professional Workstation (B). Areas within dashed lines correspond to kernel and image sizes where OpenCL is faster than threaded CPU.

**Fig. S3.**
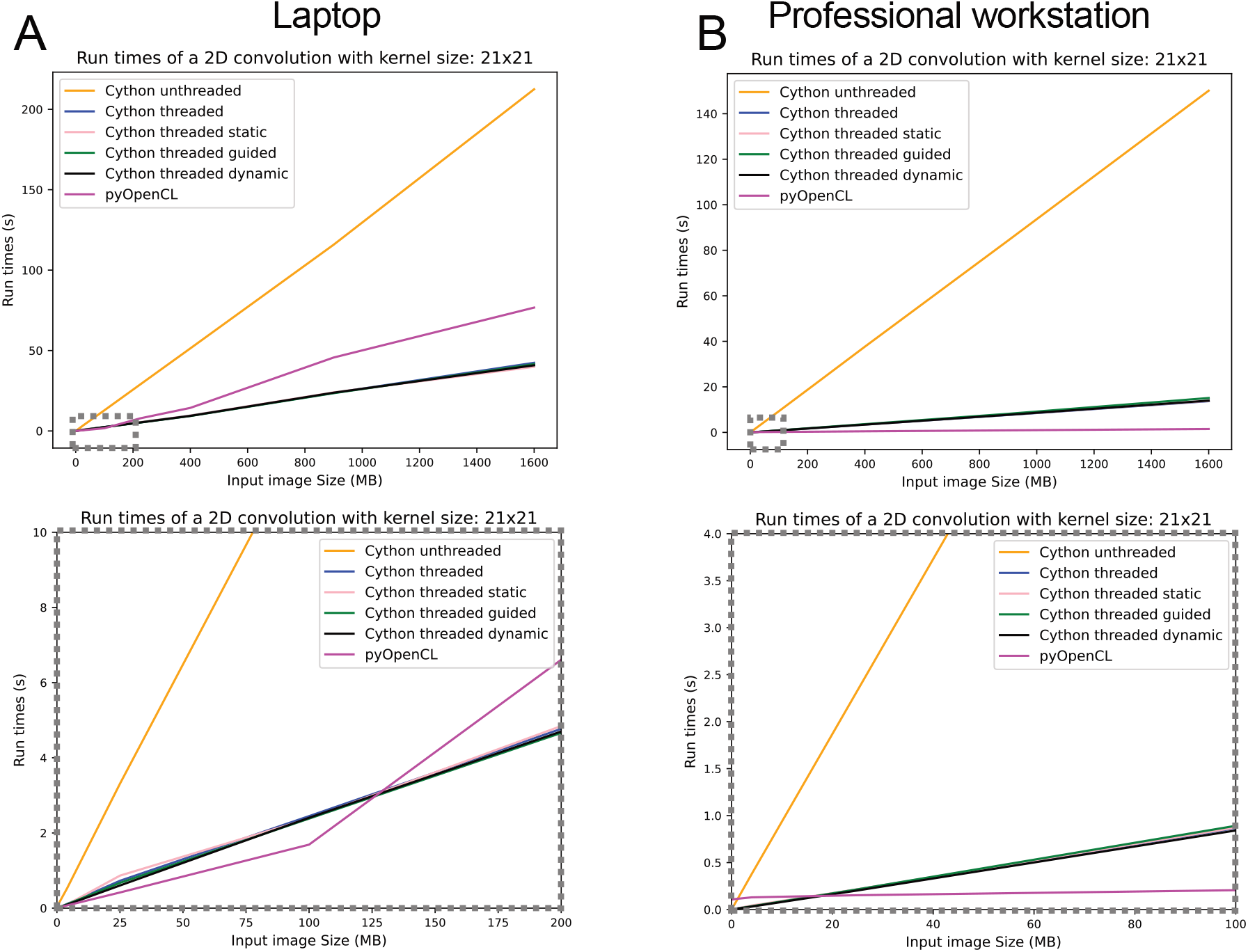
Run time of each implementation is highly dependent on the shape of input data. A 2D convolution was performed on images with increasing size using either a MacBook Air M1 (A) or a professional workstation (B). A 21 by 21 kernel was used in all operations. When using the MacBook laptop, interestingly the PyOpenCL implementation is the fastest until 125MB after which the Cython threaded implementations become significantly faster. In the professional workstation, while unthreaded is virtually always the slowest implementation, the threaded implementations are only the fastest until the size increases to 20MB, after which PyOpenCL becomes the fastest.

**Fig. S4.**
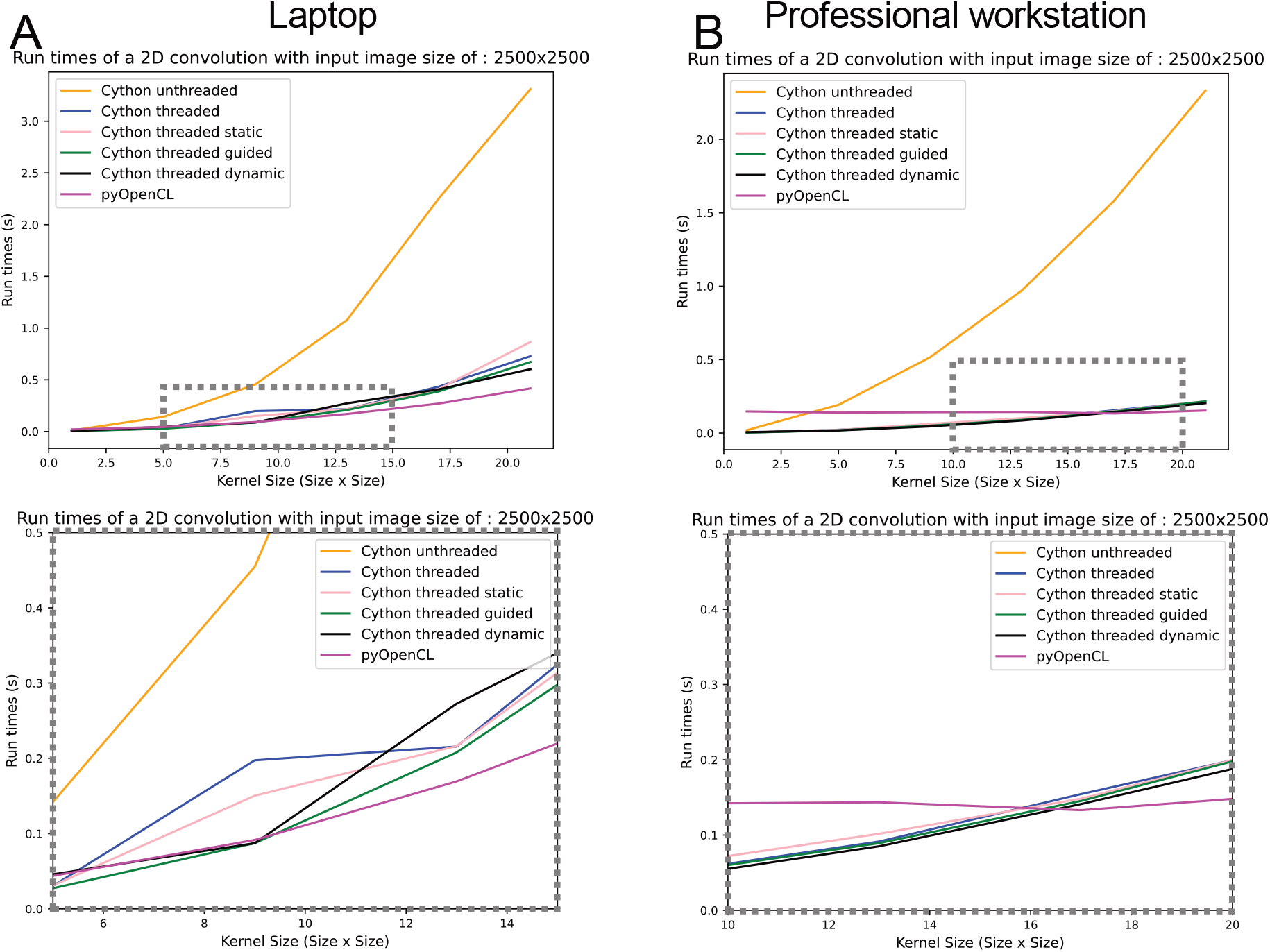
Kernel size impacts which implementation is the fastest. A 2D convolution was performed on images with varying kernel sizes, ranging from 1 to 21 (every 4) using either a MacBook Air M1 (A) or a professional workstation (B). A 21 by 21 kernel was used in all operations While unthreaded is virtually always the slowest implementation, the threaded implementations are only the fastest until the size increases to 20MB, after which PyOpenCL becomes the fastest. Bottom panels correspond to zoomed in windows of top panels, indicated by dotted boxes.

**Fig. S5.**
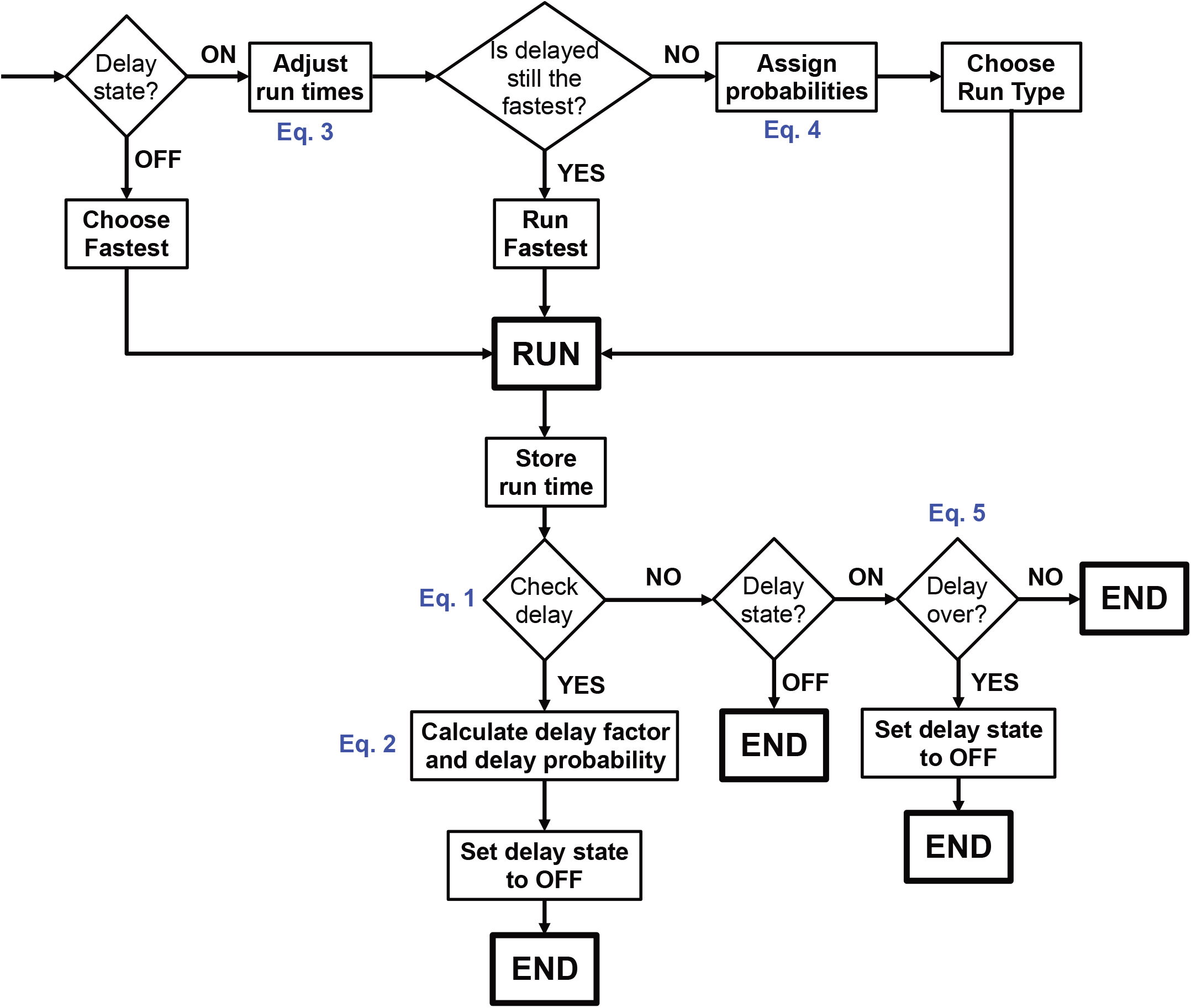
Schematic of the agent decision making for delay management.

**Fig. S6.**
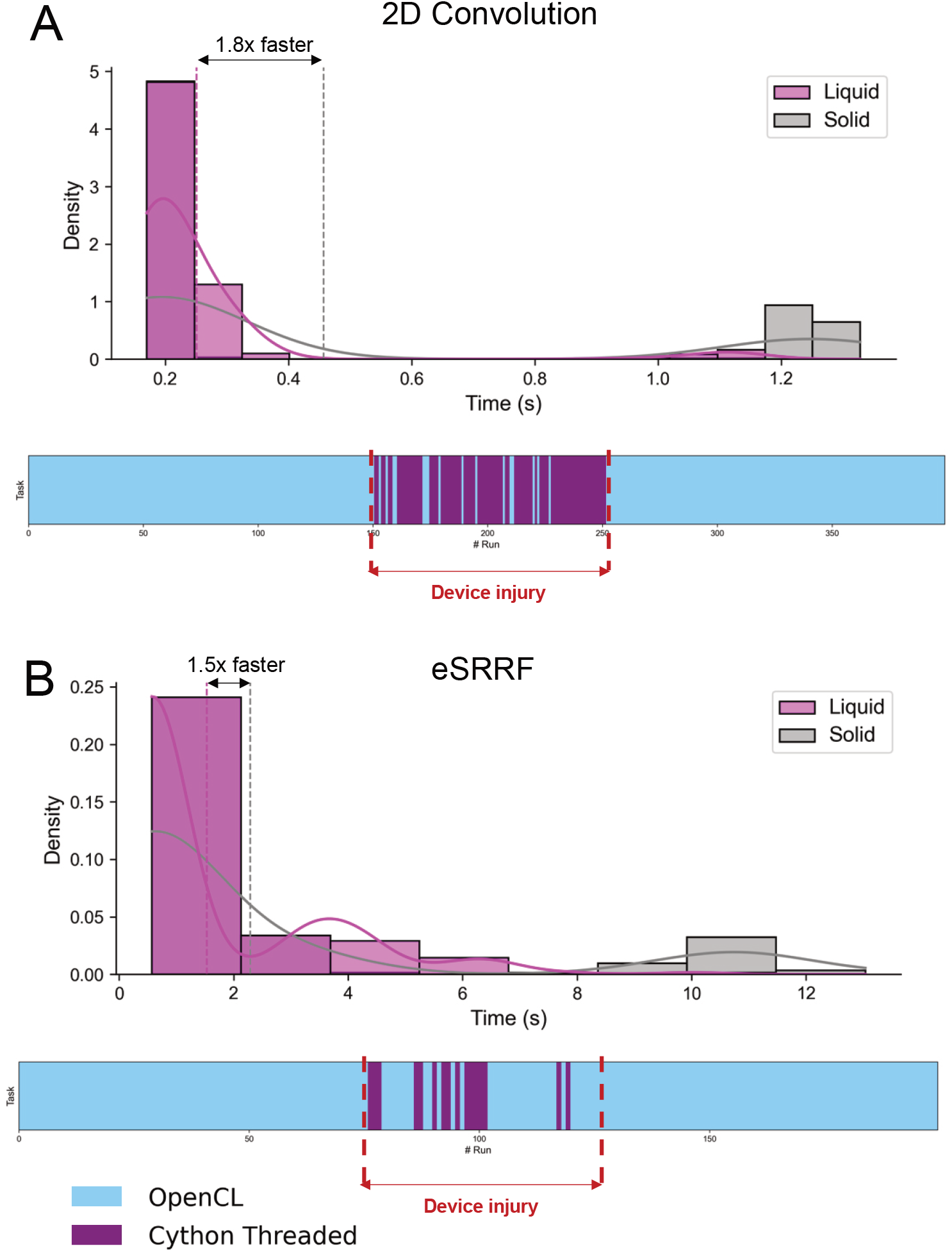
Example of delay management by the Liquid engine. Multiple two-dimensional convolutions (A) and eSRRF analysis (B) were run sequentially in a professional workstation. Starting from two initial benchmarks, the agent is responsible to inform the Liquid Engine is what is the best probable implementation. An artificial delay was induced by overloading the GPU with superfluous calculations in a separate Python interpreter.

**Fig. S7.**
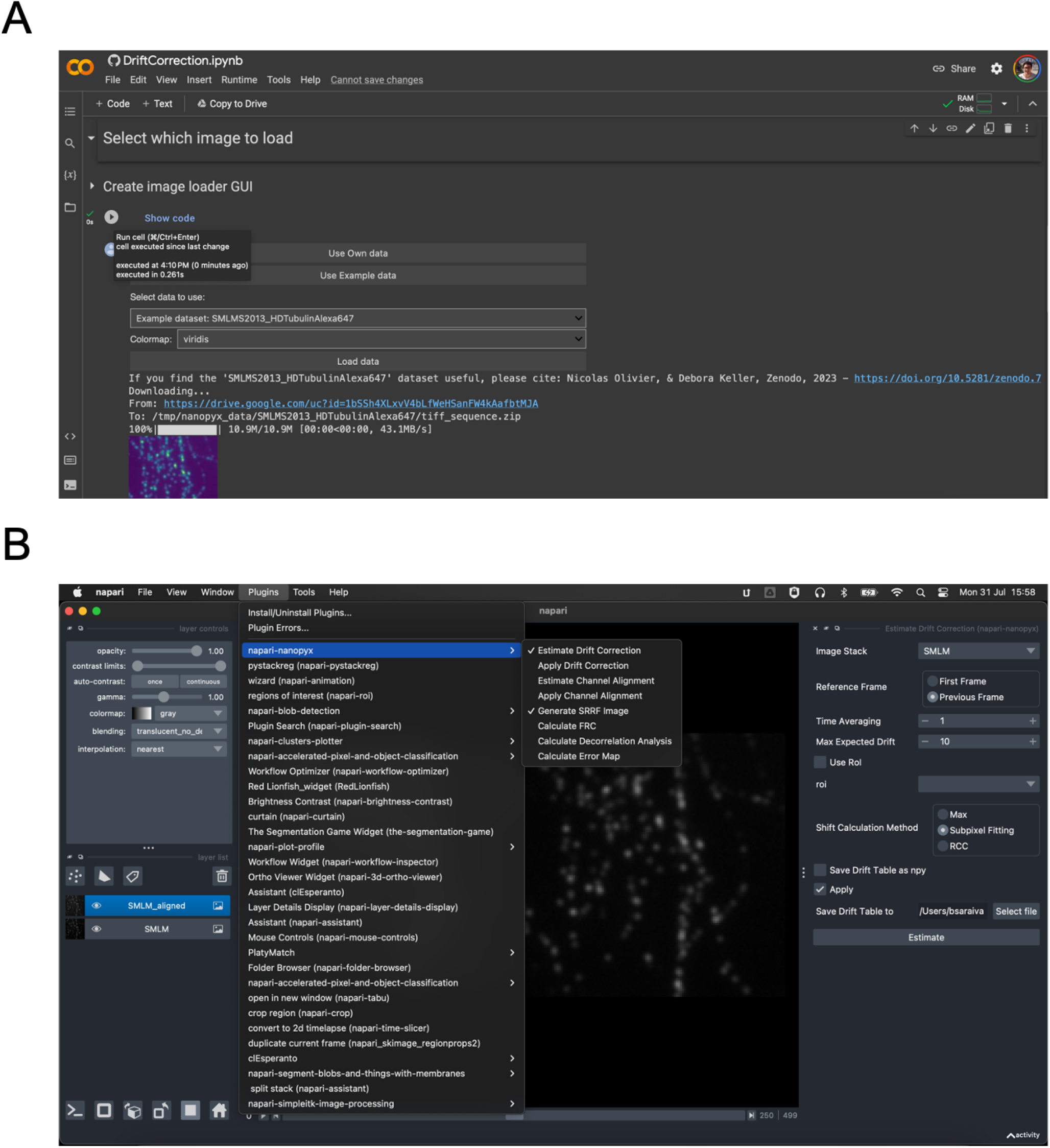
NanoPyx is available to users independently or their coding expertise. Besides the using NanoPyx as a Python library, users also have access to Jupyter notebooks (10) (A) that can either be run locally or through Google Collaboratory and a napari (11) plugin (B).

## Notes

https://github.com/HenriquesLab/NanoPyx

### Competing Interest Statement

The authors have declared no competing interest.

### Summary of Updates

Corrected incorrect acknowledgement, changing "European Commission" to "European Union".

